# Mitochondrial Redox Adaptations Enable Aspartate Synthesis in SDH-deficient Cells

**DOI:** 10.1101/2022.03.14.484352

**Authors:** Madeleine L. Hart, Evan Quon, Anna-Lena B. G. Vigil, Ian A. Engstrom, Oliver J. Newsom, Kristian Davidsen, Pia Hoellerbauer, Samantha M. Carlisle, Lucas B. Sullivan

## Abstract

The oxidative tricarboxylic acid (TCA) cycle is a central mitochondrial pathway integrating catabolic conversions of NAD+ to NADH and anabolic production of aspartate, a key amino acid for cell proliferation. Several TCA cycle components are implicated in tumorigenesis, including loss of function mutations in subunits of succinate dehydrogenase (SDH), also known as complex II of the electron transport chain (ETC). Mechanistic understanding of how proliferating cells tolerate the metabolic defects of SDH loss is still lacking. Here, we identify that SDH supports cell proliferation through aspartate synthesis but, unlike other ETC impairments, is not restored by electron acceptor supplementation. Interestingly, we find aspartate production and cell proliferation are restored to SDH impaired cells by concomitant inhibition of ETC complex I (CI). We determine that the benefits of CI inhibition in this context are dependent on decreasing mitochondrial NAD+/NADH, which drives SDH-independent aspartate production. We also find that genetic loss or restoration of SDH selects for cells with concordant CI activity, establishing distinct modalities of mitochondrial metabolism for maintaining aspartate synthesis. Collectively, these data identify a metabolically beneficial mechanism for CI loss in proliferating cells and reveal that compartmentalized redox changes can impact cellular fitness.

## Introduction

Cell proliferation requires metabolic alterations to coordinate catabolism, the maintenance of cellular bioenergetics, with anabolism, the synthesis of macromolecules for cellular replication. Several metabolic pathways serve amphibolic roles, generating both bioenergetic molecules to support metabolic homeostasis, like ATP, NADH, or NADPH, while also producing metabolic precursors used for biosynthesis of molecules such as nucleotides, amino acids, or lipids. Since these metabolic outputs are interconnected, coordinating metabolic pathway activities is critical for efficient cell proliferation. However, specific mechanisms for how metabolic alterations support cell proliferation remain poorly understood, especially in disease contexts where cell lineage, genotype, and environmental factors can all influence metabolic requirements (Vander Heiden & DeBerardinis, 2017).

Central to cell metabolism is the mitochondrion, a double membrane bound organelle that plays critical roles in cell function. While mitochondria are often viewed through the catabolic lens of efficient ATP synthesis, many proliferating cells can meet their ATP demands from aerobic glycolysis alone yet require intact mitochondrial metabolism for cell proliferation (Weinhouse et al., 1956; Tan et al., 2015; King & Attardi, 1989). Two fundamental processes of mitochondrial metabolism are the tricarboxylic acid cycle (TCA) and the electron transport chain (ETC). The TCA cycle is an amphibolic pathway, serving as both the major catabolic source of mitochondrial NADH and as a primary anabolic route supporting aspartate biosynthesis. Metabolite progression through the TCA cycle is linked to ETC activity through two mechanisms: 1) the shared succinate dehydrogenase (SDH, also known as complex II) reaction, and 2) redox coupling, where complex I (CI) accepts electrons from mitochondrial NADH to yield NAD+, in turn driving oxidative TCA cycle reactions that convert NAD+ to NADH. Disruptions to ETC complexes I, III, and IV diminish mitochondrial NAD+ regeneration, reducing the cellular NAD+/NADH ratio, impairing the TCA cycle, and slowing cell proliferation. Supplementation with molecules that regenerate NAD+, such as pyruvate, alpha-ketobutyrate, or duroquinone, can restore the NAD+/NADH ratio and the proliferation of ETC impaired cells (Birsoy et al., 2015; Luengo et al., 2021; Sullivan et al., 2015). Similarly, heterologous expression of bacterial NADH oxidase *LbNOX* can also reestablish NADH oxidation independent of the ETC and restore the proliferation of ETC impaired cells (Titov et al., 2016). These findings highlight that redox regulation is an essential function of ETC activity in proliferating cells.

A major metabolic consequence of decreased NAD+/NADH upon ETC impairment is depletion of the amino acid aspartate (Birsoy et al., 2015; Sullivan et al., 2015). Most cells are dependent on *de novo* aspartate synthesis since aspartate is poorly permeable at physiological concentrations (Garcia-Bermudez et al., 2018; Sullivan et al., 2018). Aspartate production occurs through transamination of the TCA cycle metabolite oxaloacetate, which itself is typically produced by NAD+ dependent reactions in the TCA cycle, linking aspartate production to NAD+ regeneration by the ETC. Aspartate is a central metabolic node in proliferative metabolism, serving as a direct substrate for protein synthesis and as a precursor for the synthesis of other essential metabolites, including asparagine, arginine, and both purine and pyrimidine nucleobases. Thus, aspartate depletion likely impairs core metabolic processes necessary for cell proliferation. Indeed, providing cells with exogenous sources of aspartate circumvents the proliferative defects of impairments to complexes I, III, and IV. Importantly, aspartate rescue does not correct the NAD+/NADH imbalance caused by mitochondrial inhibitors, indicating that ETC impairments block cell proliferation primarily by suppressing aspartate synthesis (Birsoy et al., 2015; Sullivan et al., 2015). Nonetheless, it remains unclear how alterations to other components of mitochondrial metabolism affect redox homeostasis and aspartate production, as well as how cells with mitochondrial dysfunction may adapt to the disruption of canonical aspartate production pathways.

The dual ETC/TCA cycle enzyme SDH serves important functions in mitochondrial metabolism yet is also a target of biological disruption. SDH is a heterotetrameric nuclear-encoded protein complex located in the inner mitochondrial membrane that catalyzes the oxidation of succinate to fumarate and shuttles electrons to ubiquinone in the ETC. Biallelic loss-of-function mutations to the subunits of SDH (A-D) or SDH assembly factors (SDHAF1-2), can lead to neoplasms in neuroendocrine tissues and the kidney (Astuti et al., 2001; Bardella et al., 2011; Baysal et al., 2000; Burnichon et al., 2010; Hao et al., 2009; Niemann & Müller, 2000). Cancer cells arising from mutations in SDH components display a complete loss of oxidative TCA cycle metabolism at the SDH step and have a substantial accumulation of the upstream metabolite succinate, which can impair enzymes involved in oxygen sensing, epigenetic regulation, and metabolism (Letouzé et al., 2013; Sullivan et al., 2016). SDH loss also blocks the canonical route for aspartate biosynthesis, making it unclear how SDH-deficient cells adapt to fulfill their anabolic demands for cell proliferation. Understanding how SDH-deficient cancer cells reconfigure their metabolism to support cell proliferation will help reveal the normal metabolic roles of SDH and could support the development of novel therapies targeting the unique metabolic liabilities of SDH-mutated cancers.

Here we investigate the consequences of acute and chronic SDH deficiency and observe that SDH impairment causes aspartate-dependent proliferation defects. However, unlike impairments to other ETC complexes, exogenous electron acceptors are insufficient to restore aspartate levels and cell proliferation to SDH-deficient cells. Surprisingly, we find that additional impairment to ETC CI is required to drive alternative aspartate synthesis in SDH-deficient cells, enabling cell proliferation. CI inhibition decreases mitochondrial NAD+/NADH, which is required to induce aspartate production and cell proliferation in SDH-impaired cells. Finally, we observe that disrupting or restoring SDH prompts selective pressure for progressive changes in CI activity to directly correspond to SDH activity, identifying distinct metabolic states that enable aspartate production. Altogether, our data reveal a novel metabolic relationship where compartmentalized redox changes are required to enact alternative aspartate biosynthetic pathways that support cell proliferation during TCA cycle dysfunction.

## Results

### SDH inhibition blocks cell proliferation, which is restored by aspartate but not electron acceptors

To understand the metabolic contributions of SDH/complex II (hereafter, SDH) to cell proliferation, we compared the proliferative consequences of the inhibitors atpenin A5 (AA5), a specific inhibitor of SDH, and the classic complex I (CI) inhibitor rotenone in the respiration-intact osteosarcoma cell line 143B. Both inhibitors effectively blocked cell proliferation in untreated media, consistent with an essential role of mitochondrial metabolism in support of cell proliferation (Figure 1A). Previous work from us and others identified that supplementation with exogenous electron acceptors pyruvate (PYR) or alpha-ketobutyrate (AKB) can restore proliferation to cells with CI impairments by regenerating NAD+ to support aspartate biosynthesis (Birsoy et al., 2015; Sullivan et al., 2015). While we confirm that PYR and AKB can robustly restore proliferation upon rotenone treatment, proliferation of AA5 treated cells was only modestly improved by electron acceptor supplementation (Figure 1A). In addition, cytosolic NAD+ regeneration by expression of *LbNOX* also restored proliferation to cells treated with rotenone, but minimally improved proliferation of cells treated with AA5 (Figures 1A and S1A). In contrast, aspartate supplementation equivalently, albeit incompletely, restored cell proliferation to both rotenone and AA5 treated cells (Figure 1A). Aspartate is poorly permeable to most cells even at supraphysiological concentrations, so we generated 143B cells expressing the glial-specific aspartate transporter SLC1A3, which we confirm supports aspartate uptake at micromolar extracellular concentrations (Figures 1B, S1B-C). Notably, improved aspartate uptake with SLC1A3 expression allowed aspartate to fully restore proliferation upon AA5 treatment, regardless of PYR co-treatment (Figure 1C). Similarly, orthogonal aspartate acquisition by intracellular expression of the guinea pig asparaginase (gpASNase1) combined with asparagine treatment was also able to restore proliferation to AA5 treated cells (Figure S1D-F) (Sullivan et al., 2018). These data indicate that aspartate is a primary metabolic limitation of both CI and SDH impaired cells, consistent with previous findings (Cardaci et al., 2015; Lussey-Lepoutre et al., 2015). However, these data also indicate that the metabolic determinants of aspartate limitation upon SDH inhibition are distinct.

**Figure 1.**
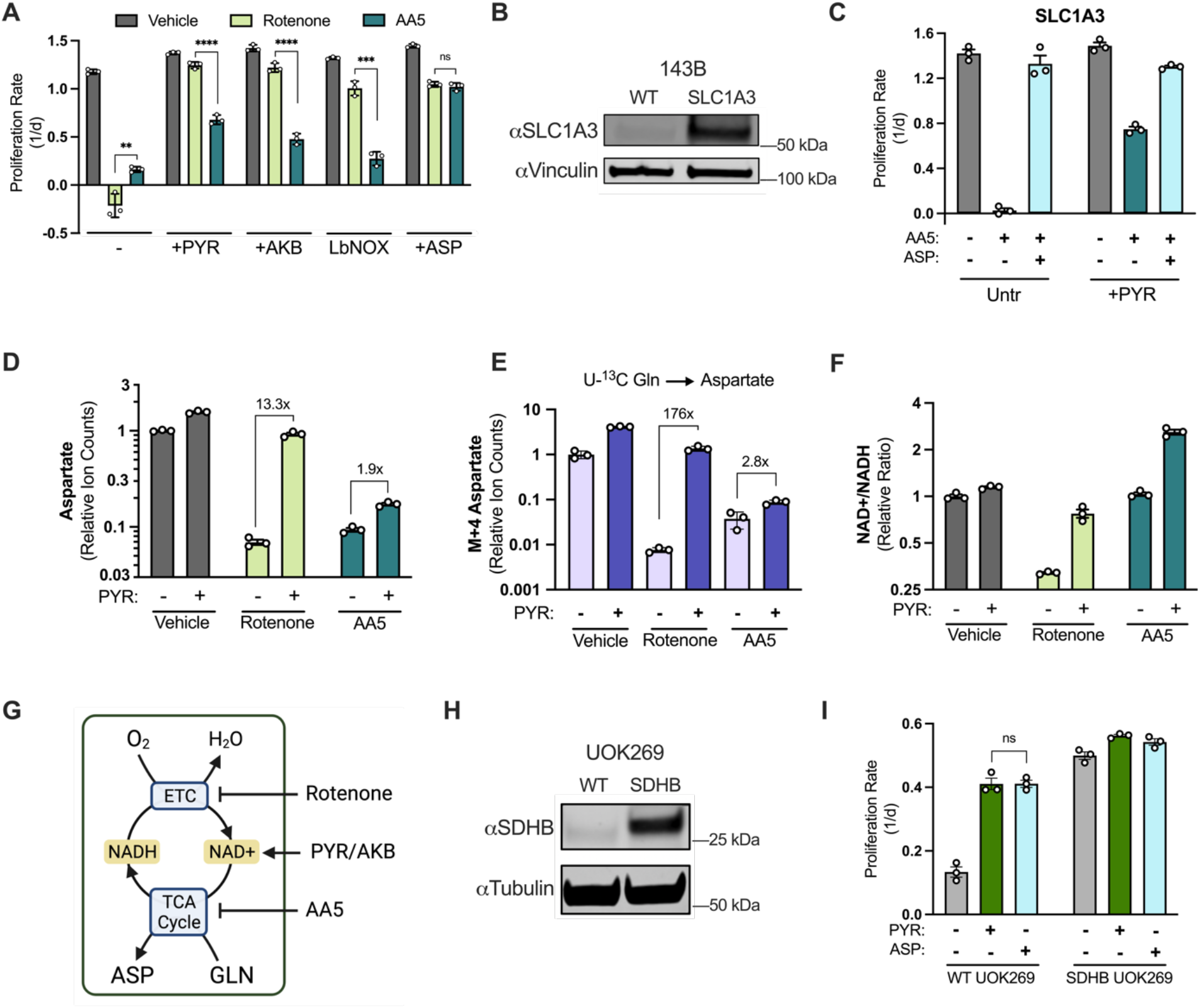
SDH inhibition blocks proliferation, which is restored by aspartate but not electron acceptors. A) Proliferation rates of 143B cells treated with vehicle (DMSO), 50 nM Rotenone, or 5 µM atpenin A5 (AA5) cultured in pyruvate free DMEM with no addition, 1 mM pyruvate (PYR), 1 mM alpha-ketobutyrate (AKB), or 20 mM aspartate (ASP). *LbNOX* expressing cells were cultured in pyruvate free DMEM without any addition (n=3). B) Western blot for SLC1A3 and vinculin in 143B cells expressing SLC1A3 or wild type (WT) control 143B cells. Vinculin is used as a loading control. C) Proliferation rates of SLC1A3 expressing 143B cells treated with vehicle (DMSO), 5 µM AA5 or 5 µM AA5 and 1 mM aspartate in pyruvate free DMEM with no addition or 1 mM PYR (n=3). D) Aspartate levels measured by liquid chromatography-mass spectrometry (LCMS) metabolomics in 143B cells treated with vehicle (DMSO), 50 nM rotenone, or 5 µM AA5 in pyruvate free DMEM with no addition or 1 mM PYR (n=3). E) M+4 aspartate levels measured by LCMS metabolomics from 143B cells cultured with pyruvate/glutamine free DMEM supplemented with 4 mM U-^13^C glutamine and treated with vehicle (DMSO), 50 nM rotenone, or 5 µM AA5 in the absence and presence of 1 mM PYR for 6 hours (n=3). F) The NAD+/NADH ratio measured by LCMS metabolomics of 143B cells cultured in pyruvate free DMEM and treated with vehicle, 50 nM rotenone, or 5 µM AA5 in pyruvate free DMEM with no addition or 1 mM PYR for 6 hours (n=3). G) Schematic showing that rotenone inhibits the ETC, blocking NADH oxidation causing an indirect TCA cycle impairment that can be overcome by exogenous electron acceptors (PYR/AKB), while AA5 directly blocks the TCA cycle and aspartate synthesis. H) Western blot for SDHB and α-tubulin (tubulin) in WT UOK269 cells or UOK269 cells with restoration of SDHB (SDHB UOK269) as indicated. Tubulin is used as a loading control. I) Proliferation of WT UOK269 and SDHB UOK269 cells cultured in pyruvate free DMEM supplemented with vehicle (H_2_O), 1 mM PYR, or 20 mM ASP (n=3). Fold change is indicated above brackets in D and E. Data are plotted as means ± standard deviation (SD) and compared with an unpaired two-tailed student’s t-test. p<0.05*, p<0.01**, p<0.001***, p<0.0001****

To evaluate the metabolic differences between impairments to CI or SDH, we conducted liquid chromatography-mass spectrometry (LCMS) metabolomics on cells treated with either inhibitor, with or without electron acceptor supplementation. As expected for SDH impairment, AA5 treatment caused succinate accumulation and depleted fumarate levels (Figure S1G-H). Consistent with aspartate being the mediator of proliferation defects from either inhibitor, aspartate levels were depleted by both drugs, fully restored by pyruvate in rotenone treated cells, and only partially increased in cells treated with AA5 (Figure 1D). To ascertain the source of aspartate production in each condition we measured the generation of M+4 aspartate in cells cultured in U-^13^C glutamine, reflective of aspartate produced via canonical oxidative TCA cycle activity, and found that pyruvate treatment restores metabolic progression through the TCA cycle in rotenone treated cells but not in cells treated with AA5 (Figure 1E). Unlike with rotenone treatment, AA5 treatment was not associated with suppression of NAD+/NADH, breaking the direct correlation between NAD+/NADH and aspartate levels seen with other ETC inhibitors (Figure 1F) (Gui et al., 2016). Pyruvate supplementation increased NAD+/NADH in both cases; restoring NAD+/NADH in rotenone treated cells to near vehicle treated levels and causing an NAD+/NADH increase beyond vehicle treated cells upon AA5 treatment (Figure 1F). Collectively, these data are consistent with a model where CI impairments suppress aspartate production by lowering NAD+/NADH and thereby slowing the oxidative TCA cycle, whereas SDH inhibition directly blocks the oxidative TCA cycle, impairing aspartate and NADH production (Figure 1G).

We also evaluated the metabolic effects of altered SDH activity in UOK269 cells, a patient derived renal cell carcinoma cell line that arose with biallelic loss of function mutations in SDHB (Saxena et al., 2015). We re-expressed wild type SDHB in UOK269 cells, which alleviated the accumulation of succinate and permitted oxidative TCA cycle-derived aspartate from U-^13^C glutamine (Figures 1H, S1I-K). Similar to 143B cells, intact TCA cycle activity ameliorated the dependence of UOK269 cells on pyruvate or aspartate for cell proliferation (Figure 1I). However, in contrast to AA5 treated 143B cells, pyruvate and aspartate treatment were equally effective at restoring cell proliferation in UOK269 cells (Figure 1I). These data suggest that long-term SDH loss may be associated with metabolic adaptations that allow cells to fully take advantage of exogenous electron acceptors to restore aspartate and cell proliferation.

### SDH-deficient cells require CI inhibition for aspartate synthesis and cell proliferation

We next questioned what metabolic adaptations may occur in SDH-deficient cells to promote cell proliferation. Intriguingly, several cell lines generated with genetic defects in SDH components have diminished respiration rates (Cardaci et al., 2015; Lorendeau et al., 2017; Saxena et al., 2015). Since CI and SDH are functionally distinct contributors to the ETC, and CI derived electrons account for the majority of respiration, it is not inherently obvious that SDH-deficient cells would be unable to maintain robust respiration activity. In support of this, we found that treatment with AA5 only partially impairs respiration after several hours of treatment, whereas rotenone severely abrogates respiration within minutes (Figure S2A). One potential explanation for the respiration loss in SDH-mutant cells is the observation that they manifest with decreased CI expression and activity (Cardaci et al., 2015; Lorendeau et al., 2017). Indeed, dual inhibition of SDH and CI was reported to be required to mimic the metabolic phenotype of SDH-mutant cancer cells (Lorendeau et al., 2017). Nonetheless, while these data suggest that CI loss may serve a metabolic role in SDH-deficient cells, no functional benefit for CI loss in this context has yet been described.

We tested if, in the presence of electron acceptors, rotenone could alter the proliferation of cells treated with AA5 and, surprisingly, found it caused a near complete proliferation restoration (Figure 2A). We then measured aspartate levels and found that CI co-inhibition partially restored aspartate levels to SDH-impaired cells (Figure 2B). CI co-inhibition also restored proliferation and aspartate levels across a panel of diverse human cell lines treated with AA5, including both transformed (A549, HCT116) and non-transformed (293T, TF-1) cells, indicating that this metabolic relationship was generalizable (Figure S2B, S2C). To test these effects genetically, we used CRISPR/Cas9 to generate gene knockouts (KO) of SDHB, the most frequently mutated subunit in SDH-null human cancers (Amar et al., 2007; Badenhop et al., 2004; Klein et al., 2008; Linehan & Ricketts, 2013). While we were able to achieve efficient disruption of SDHB, with ∼96% loss in 143B cells, we were unable to generate single cell SDHB KO clones in standard media (Figure 2C). Based on our previous findings, we repeated colony selection with or without rotenone and found that CI inhibition was essential for successful generation of clones from sgSDHB nucleofected 143B cells (Figure 2D). Notably, every tested clone generated in this manner was devoid of SDHB expression (Figures 2E and S2D). In each SDHB KO cell line generated, proliferation was slowed relative to their parental cells but was restored by treatment with the CI inhibitors rotenone or metformin, or by supplementation with aspartate (Figures 2F, S2E). We also restored SDHB expression in a KO clone and found that it rescued proliferation to parental rates and reinstated wild type relationships with rotenone and AA5 treatment (Figures S2F, S2G). We then measured aspartate levels in WT and SDHB KO cells with or without rotenone treatment and found that rotenone decreases aspartate levels in WT cells but increases aspartate in SDHB KO cells, consistent with proliferation restoration (Figure 2G). Finally, to test if this metabolic relationship was an artifact of our cell culture media choice, we also tested the effects of SDH inhibition in 143B cells cultured in Human Plasma-Like Media (HPLM), which closely matches the nutrient composition of human plasma (Cantor et al., 2017). Importantly, we found that AA5 still impaired cell proliferation and that rotenone or aspartate co-treatment restored cell proliferation in HPLM (Figure S2H). Collectively, our data demonstrate that CI loss is required for the aspartate-dependent proliferation in diverse cell lines and media conditions upon SDH disruption.

**Figure 2.**
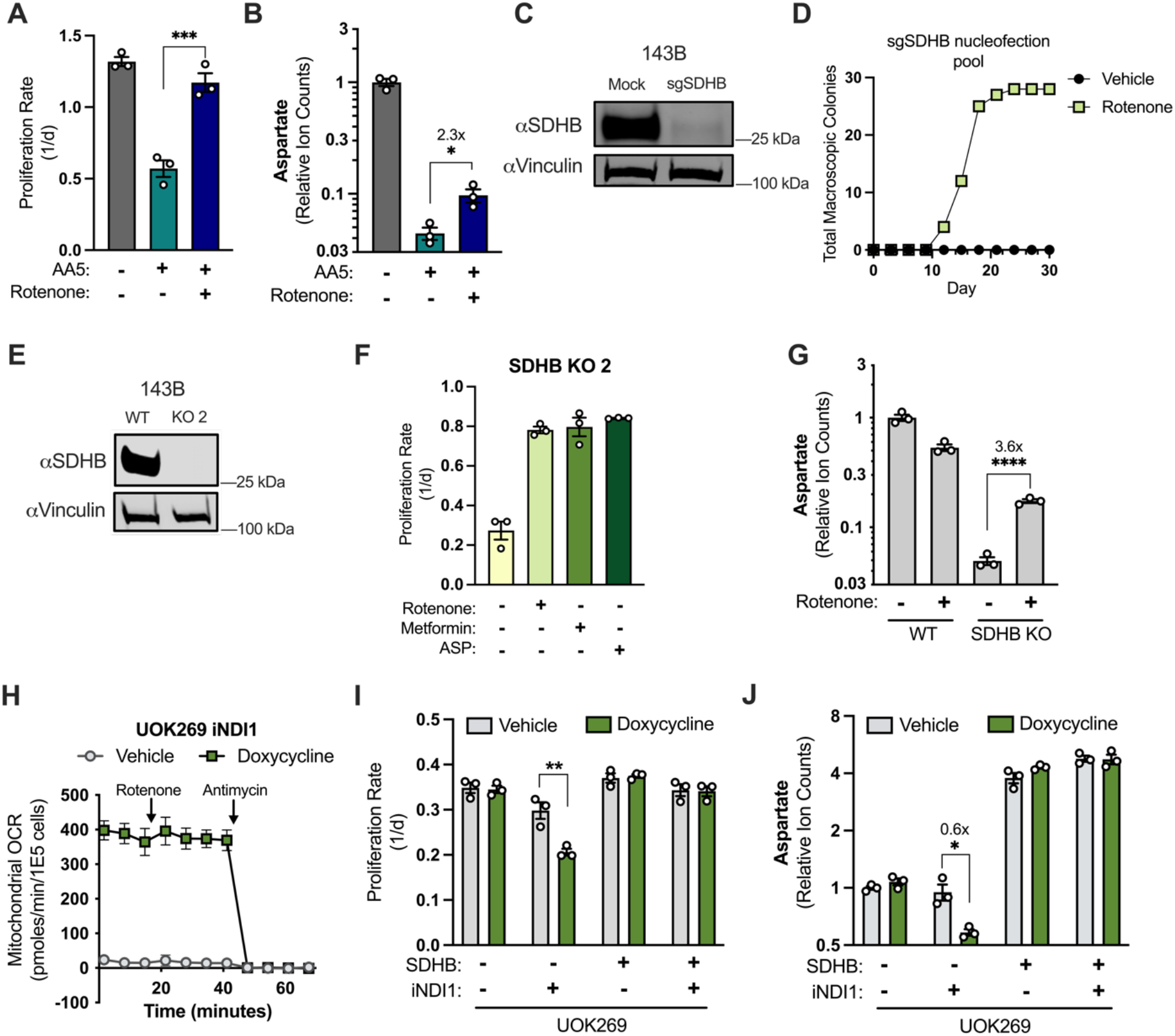
SDH-deficient cancer cells require CI inhibition for aspartate synthesis and proliferation. A) Proliferation rates of 143B cells cultured in DMEM treated with vehicle (DMSO), 5 µM AA5, or 5 µM AA5 and 50 nM rotenone (n=3). B) Aspartate levels of 143B cells cultured in DMEM treated with vehicle (DMSO), 5 µM AA5, or 5 µM AA5 and 50 nM rotenone for 6 hours (n=3). C) Western blot for SDHB and vinculin from 143B cells 3 days after nucleofection with plasmid GFP or sgRNAs and sNLS-SpCas9 for SDHB as indicated. Vinculin is used as a loading control. D) Number of single cell clones that formed colonies from sgSDHB pool in 143B cells treated with vehicle (DMSO) or 30 nM rotenone during a 30 day period. E) Western blot for SDHB and vinculin from WT 143B cells and SDHB KO 143B clone 2. Vinculin is used as a loading control. F) Proliferation rates of SDHB KO 143B cells (clone 2) cultured in DMEM and treated with vehicle (DMSO), 50 nM rotenone, 1 mM metformin, or 20 mM ASP (n=3). G) Aspartate levels measured by LCMS metabolomics of WT 143B cells and SDHB KO 2 cells cultured in DMEM and treated with vehicle (DMSO) or 50 nM rotenone for 6 hours (n=3). H) Mitochondrial oxygen consumption rates of pInducer20-NDI expressing UOK269 cells (UOK269 iNDI1) that were pre-treated with vehicle (H_2_O) or 1 µg/mL doxycycline for 40 hours as indicated, followed by injections of 100 nM rotenone and 10 µM antimycin (n=3). I) Proliferation rates of WT and SDHB UOK269 cells with or without expression of pInducer20-NDI1 (iNDI1) cultured in DMEM and treated with vehicle (H_2_O) or 1 µg/mL doxycycline (n=3). J) Aspartate levels measured by LCMS metabolomics of WT and SDHB UOK269 cells with or without expression of pInducer20-NDI1 (iNDI1) cultured in DMEM and treated with vehicle (H_2_O) or 1 µg/mL doxycycline for 24 hours (n=3). Fold change is indicated above brackets in B, G and J. Data are plotted as means ± standard deviation (SD) except in H which are means ± standard error of the mean (SEM) and compared with an unpaired two-tailed student’s t-test. p<0.05*, p<0.01**, p<0.001***, p<0.0001****

One potential caveat with generalizing these effects to SDH mutant cancers is that SDH wild type parental cell lines may not recapitulate the intrinsic metabolic features of naturally arising SDH-mutant cancer cells. We measured the CI activity of the SDH-null UOK269 cells and found substantially decreased CI activity compared to WT 143B cells (Figure S2I). We therefore hypothesized that if CI restoration would modify aspartate production and cell proliferation in UOK269 cells. Mammalian CI activity is dependent on the expression of >50 genes encoding core subunits and assembly factors, so increasing CI activity by traditional gene overexpression is not feasible. Instead, we expressed a doxycycline-inducible construct of the *Saccharomyces Cerevisiae* NDI1, a rotenone-insensitive CI analog that can restore NADH oxidation and ETC activity in mammalian cells when CI is impaired (Seo et al., 1998; Wheaton et al., 2014). Importantly, doxycycline treatment of wild type UOK269 cells with the inducible NDI1 (iNDI1) construct resulted in a substantial induction of mitochondrial oxygen consumption that was resistant to rotenone treatment, verifying NDI1 expression and confirming that decreased CI activity is a bottleneck for ETC function in UOK269 cells (Figure 2H). While NDI1 expression had no effect on the proliferation of SDHB-restored UOK269 cells, NDI1 induction specifically decreased the proliferation of wild type SDH-null UOK269 cells (Figure 2I). Metabolomics measurements also found that NDI1 expression specifically lowered aspartate levels in wild type UOK269 cells, but not in their SDHB-restored counterparts (Figure 2J).

### Mitochondrial redox alterations are essential for CI inhibition to rescue SDH-deficient cells

We next investigated the metabolic mechanism for how CI impairment could be providing a benefit to SDH-deficient cells. CI activity contributes to several metabolic processes, including NADH oxidation, mitochondrial membrane potential generation, and the reduction of ubiquinone (Figure S3A). To determine which of these processes is deleterious to SDH-deficient cells, we compared co-treatment of AA5 with rotenone or oligomycin, two inhibitors with distinct mechanisms of action for inhibiting ETC activity. While rotenone directly inhibits CI, oligomycin blocks the mitochondrial inner membrane complex ATP synthase, inhibiting the consumption of the mitochondrial membrane potential, inducing mitochondrial inner membrane hyperpolarization, causing collateral inhibition of ETC activity. Oligomycin treatment thereby similarly impairs NADH oxidation but differs from CI inhibitors by increasing the mitochondrial membrane potential and slowing electron flux through all three proton pumping complexes (Figure S3A). Importantly, treatment with oligomycin can substitute for CI inhibitors, albeit less effectively, to restore the proliferation and aspartate levels of AA5 treated cells (Figure S3B-C). These results suggest that altered mitochondrial NAD+/NADH, rather than other downstream effects on ETC function, is likely a primary mechanism by which CI inhibition supports aspartate production and proliferation in SDH-deficient cells.

To determine the effects of CI inhibition on the redox state of SDH-impaired cells, we measured NAD+/NADH by LCMS. While AA5 treatment alone increased NAD+/NADH (Figure 1F), the effects of rotenone were epistatic to AA5 treatment and reduced the whole cell NAD+/NADH when co-treated (Figure 3A). Since the proximate effect of rotenone treatment on NAD+/NADH occurs in the mitochondria and these cells can maintain cytosolic NAD+/NADH using environmental pyruvate (Williamson et al., 1967), we hypothesized that mitochondrial localized redox changes may drive the proliferative benefits of CI inhibition upon SDH impairment. To determine if mitochondrial NAD+/NADH alterations are required for the proliferation benefits of CI inhibition in SDH-impaired cells, we expressed cytosolic or mitochondrial-targeted FLAG-tagged *Lactobacillus brevis* NADH oxidase (cyto*LbNOX*/mito*LbNOX*) in 143B cells, which can promote NADH oxidation independent of ETC activity (Figure 3B) (Titov et al., 2016). Subcellular fractionation and immunoblotting verified successful subcellular targeting, as each was enriched in its expected compartment (Figure 3C). Whereas cells expressing cyto*LbNOX* behaved similarly to parental cells, mito*LbNOX* expression blocked the proliferation restorative effect of rotenone treatment in SDH-impaired cells (Figure 3D). Moreover, the increase in aspartate levels upon rotenone co-treatment was diminished by mito*LbNOX* expression, but not by cyto*LbNOX* expression (Figure 3E). These data indicate that decreased mitochondrial NAD+/NADH is required for the benefits of CI inhibition when SDH is impaired. We next asked if the benefits of decreased mitochondrial NAD+/NADH in SDH-impaired cells were separable from other effects of CI inhibition. Another determinant of mitochondrial NAD+/NADH is the pyruvate dehydrogenase (PDH) reaction, which converts NAD+ to NADH (Figure 3F). The drug AZD7545 can disinhibit PDH by blocking regulation by pyruvate dehydrogenase kinases (PDKs) (Morrell et al., 2003). In the context of electron acceptor limitation, AZD7545 can further drive down NAD+/NADH and impair cell function (Luengo et al., 2021); however, our data indicate that lowering mitochondrial NAD+/NADH is beneficial in SDH-deficient cells, so we hypothesized that AZD7545 may instead provide a benefit in this context. To ensure that the AZD7545 mechanism of action functions independently of CI inhibition, we confirmed that AZD7545 treatment does not impair mitochondrial oxygen consumption in AA5 treated cells (S3D). Additionally, AA5 treated cells co-treated with AZD7545 also maintained glucose-derived cis-aconitate levels, whereas rotenone co-treatment depleted them, highlighting the distinct metabolic mechanisms by which these inhibitors drive mitochondrial redox changes (Figure S3E). Importantly, treatment with AZD7545 also improved cell proliferation and increased aspartate levels in SDH-impaired cells (Figure 3G-H). Collectively, these data indicate that compartment-specific redox alterations are both sufficient and required to reprogram aspartate synthesis in SDH-impaired cells.

**Figure 3.**
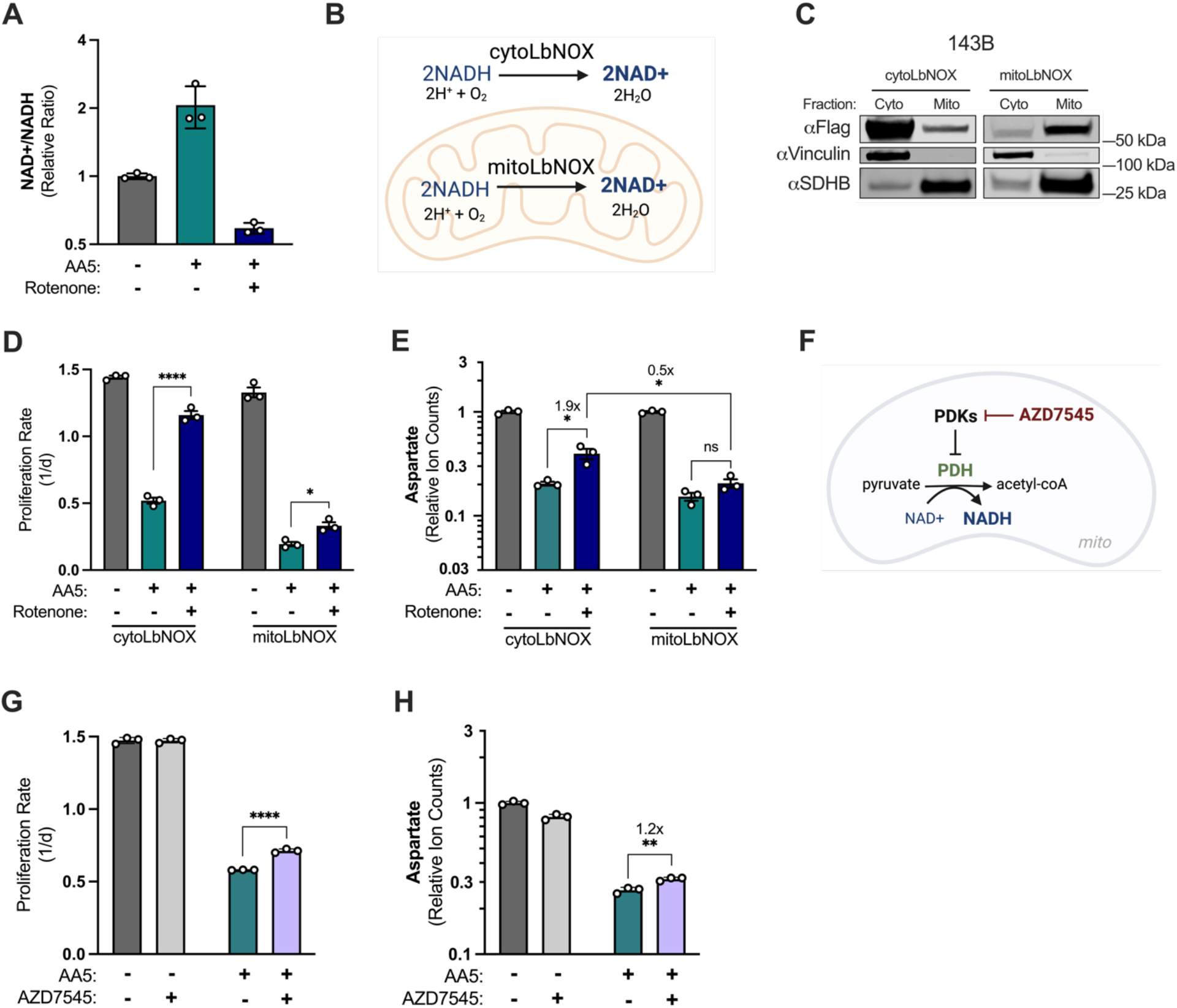
CI inhibition decreases mitochondrial NAD+/NADH, which is required for aspartate synthesis and proliferation in SDH-impaired cells. A) Whole cell NAD+/NADH ratio measured by LCMS metabolomics of 143B cells cultured in DMEM and treated with vehicle (DMSO), 5 µM AA5, or 5 µM AA5 and 50 nM rotenone for 6 hours (n=3). B) Schematic depicting the functions of cyto*LbNOX* and mito*LbNOX* is each compartment as indicated. C) Western blot for FLAG, Vinculin, and SDHB from 143B cells expressing FLAG-tagged cyto*LbNOX* or mito*LbNOX*, in cytosolic or mitochondrial fractions isolated by differential centrifugation. Vinculin is a loading control for cytosol and SDHB is a loading control for mitochondria. D) Proliferation rates of cyto*LbNOX* and mito*LbNOX* expressing 143B cells cultured in DMEM treated with vehicle (DMSO), 5 µM AA5, or 5 µM AA5 and 50 nM rotenone (n=3). E) Aspartate levels measured by LCMS metabolomics of cyto*LbNOX* and mito*LbNOX* expressing 143B cells cultured in DMEM and treated with vehicle (DMSO), 5 µM AA5, or 5 µM AA5 and 50 nM rotenone for 6 hours (n=3). F) Schematic showing the mechanism of action of AZD7545 to activate pyruvate dehydrogenase (PDH) by inhibition of the negative regulators of PDH, pyruvate dehydrogenase kinases (PDKs). G) Proliferation rates of WT 143B cells cultured in DMEM and treated with vehicle (DMSO), 5 µM AA5, 5 µM AZD7545, or 5 µM AA5 and 5 µM AZD7545 (n=3). H) Aspartate levels measured by LCMS metabolomics of WT 143B cells cultured in DMEM and treated with vehicle (DMSO), 5 µM AA5, 5 µM AZD, or 5 µM AA5 and 5 µM AZD for 6 hours (n=3). Fold change is indicated above brackets in E and H. Data are plotted as means ± standard deviation (SD) and compared with an unpaired two-tailed student’s t-test. p<0.05*, p<0.01**, p<0.001***, p<0.0001****

### SDHB-null cells adapt through progressive complex I loss

After several months of passaging SDHB KO cells, we observed that their proliferation rate increased and that the benefit of treatment with CI inhibitors or supplementing with aspartate was lost (Figure 4A). In addition to these 143B SDHB KO cells, we also generated HEK293T SDHB KO cells and found similar results (Figure S4A-B). To understand this phenomenon, we generated three categories of cells derived from an SDHB KO clone: Early Passage (EP), <5 passages; Late Passage (LP), >15 passages; or Addback (AB), where SDHB was re-expressed immediately after SDHB KO clone derivation (Figure S4C). We measured metabolite levels comparing EP and LP cells and found that aspartate was basally increased in the LP cells, consistent with their increased proliferation rates, and that the induction of aspartate upon rotenone treatment was blunted (Figure 4B). To determine if this adaptive effect was due to a selective proliferation advantage or simply inherent to passaged SDH-null cells, we separately passaged EP SDHB KO cells in rotenone for 15 passages (Figure S4D). We found that cells cultured with continuous rotenone treatment remained phenotypically similar to EP cells when CI inhibitors were removed, suggesting the selective adaptive pressure is stalled by exogenous CI inhibition (Figure S4E).

**Figure 4.**
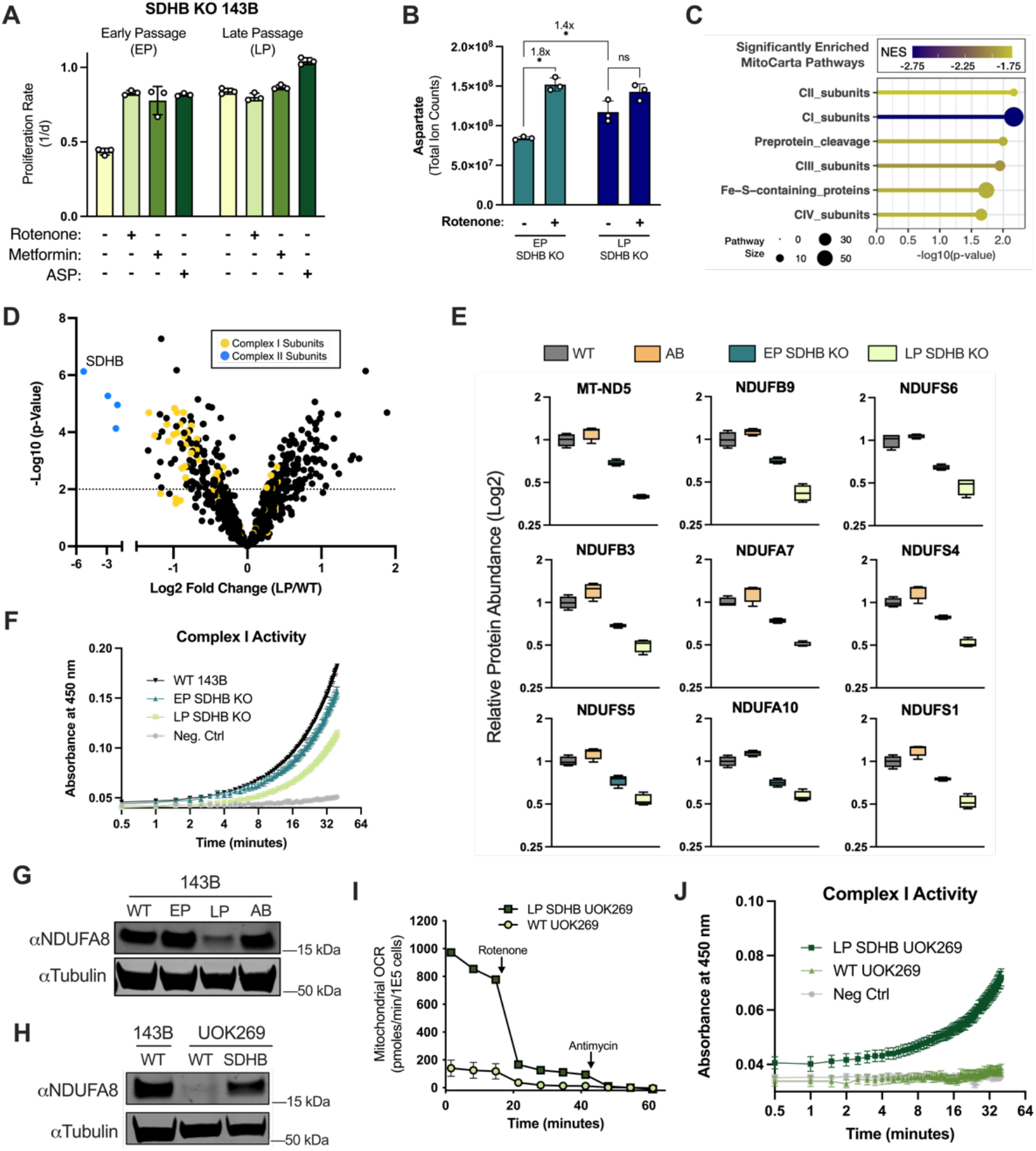
CI activity loss is a metabolic adaptation that supports proliferation in SDHB-null cells. A) Proliferation rates of early passage (EP) and late passage (LP) SDHB KO 143B cells cultured in DMEM and treated with vehicle (DMSO), 50 nM rotenone, 1 mM metformin, or 20 mM ASP (n=3). B) Aspartate levels measured by LCMS metabolomics of EP and LP SDHB KO 143B cells cultured in DMEM treated with vehicle (DMSO) or 50 nM rotenone for 6 hours (n=3). C) Gene set enrichment analysis of LP SDHB KO 143B cells compared to WT 143B cells using MitoCarta 3.0 pathways and mitochondrial proteomics data. D) Volcano plot of compiled LP SDHB KO 143B cells compared to WT 143B cells showing all mitochondrial proteins (black), complex II subunits (blue), and complex I subunits (yellow) (n=4). E) Box plots of a subset of CI subunits showing relative expression of mitochondria from WT 143B, SDHB AB 143B, EP SDHB KO 143B, and LP SDHB KO 143B cells (n=4). F) Complex I activity assay of WT 143B, EP SDHB KO, and LP SDHB KO cells (n=2). G) Western blot for NDUFA8 and tubulin from WT 143B, EP SDHB KO, LP SDHB KO, and SDHB AB cells. Tubulin is used as a loading control. H) Western blot for NDUFA8 and tubulin from WT 143B, WT UOK269, and LP SDHB UOK269 cells. Tubulin is used as a loading control. I) Mitochondrial oxygen consumption rate of WT UOK269 cells compared to LP SDHB UOK269 cells with indicated injections of 100 nM rotenone and 10 µM antimycin (n=3). J) Complex I activity assay (Abcam) of WT UOK269 and LP SDHB UOK269 cells (n=3). Fold change is indicated above brackets in B. Data are plotted as means ± standard deviation (SD) except in I and J which are means ± standard error of the mean (SEM) and compared with an unpaired two-tailed student’s t-test. p<0.05*, p<0.01**, p<0.00***, p<0.0001****

To learn more about the mitochondrial alterations occurring in SDHB KO cells, we conducted tandem mass tag (TMT) quantitative proteomics on isolated mitochondria from WT cells, AB cells, EP SDHB KO cells, and LP SDHB KO cells (Figure S4C). Unsupervised clustering of protein levels found that each group clustered together and that both pairs of SDHB KO and replete cells clustered together (Figure S4F). Comparing WT and LP samples, we conducted MitoPathways 3.0 pathway enrichment analysis (Pagliarini et al., 2008; Rath et al., 2021) of mitochondrial protein abundance changes and found that following complex II proteins, CI proteins were the most significantly enriched as different (Figure 4C). Indeed, the majority of CI subunits were decreased in abundance in LP SDHB KO cells compared to WT cells (Figure 4D). In addition, CI subunit levels progressively decreased from WT/AB to EP SDHB KO to LP SDHB KO, which resulted in decreased CI activity, further suggesting that SDHB KO cells select against CI activity over time (Figure 4E-F). We confirmed these findings by western blot, where LP SDHB KO 143B cells were associated with an ∼80% reduction in the CI subunit NDUFA8 compared to WT cells (Figure 4G). Finally, we also tested the reverse experiment, evaluating if SDHB-replenished UOK269 cells are associated with restoration of CI. Indeed, late passage SDHB UOK269 cells regained NDUFA8 expression, mitochondrial oxygen consumption, and CI activity when compared to their SDH-deficient wild type counterparts (Figure 4H-J). Together, these data indicate that CI activity is reversibly altered to generate an ideal mitochondrial redox state to enable aspartate synthesis.

## Discussion

Mitochondria are established contributors to catabolic cell metabolism through their canonical oxidative TCA cycle and ETC activities. In proliferating cells, the anabolic contributions of these processes in NAD+/NADH homeostasis and TCA cycle-derived aspartate production have recently gained appreciation, defining important amphibolic roles for oxidative mitochondrial metabolism (Birsoy et al., 2015; Sullivan et al., 2015; Titov et al., 2016). However, the fact that loss of function mutations in genes encoding the TCA cycle enzymes SDH and Fumarate Hydratase (FH) can promote tumorigenesis in some contexts challenges the notion that mitochondrial oxidative anabolism is strictly essential for aspartate production. Here, we investigate the metabolic roles of SDH and find that, in cells that employ oxidative TCA cycle function, SDH activity is primarily required for cell proliferation by supporting aspartate production. Interestingly, we found that cells can tolerate SDH loss if they also have concomitant CI impairment, which supports cell proliferation by decreasing mitochondrial NAD+/NADH to enable alternative aspartate synthesis. The linked nature of SDH and CI was also revealed by reciprocal experiments where either SDHB was knocked out in wild type cells or restored in cancer cells arising from SDHB mutations. In both cases, functional SDH selects for cells with high CI activity and impaired SDH selects for cells with low CI activity, suggesting two modalities for mitochondrial anabolic metabolism: 1) CI regenerates NAD+ to drive oxidative anabolism via the TCA cycle, or 2) CI loss prevents NADH consumption, supporting NADH-driven reductive anabolism in the mitochondria of cells without TCA cycle function. To our knowledge, this work provides the first mechanistic evidence that CI loss can directly support core metabolic functions, in this case by fueling aspartate synthesis for cell proliferation.

Metabolic changes are interconnected with tumorigenesis, which is epitomized by cancers deriving from mutations in TCA cycle genes. We found that cancer cells devoid of SDH activity enact further metabolic changes by suppressing CI to maximize cell proliferation, suggesting that these cells may have distinct metabolic requirements that could be targeted to impair tumor growth. However, cancer cell metabolic requirements can differ in animals, so evaluating these metabolic variables in complex tumor microenvironments will be essential prior to leveraging them for cancer treatment (Davidson et al., 2016). Nonetheless, the phenomenon of CI loss has also been observed in several other kidney neoplasias, including FH-mutant type 2 papillary renal cell carcinomas (pRCC), renal oncocytomas driven by mitochondrial DNA mutations, and VHL-mutant clear cell renal cell carcinomas (ccRCC), which are characterized by suppression of oxidative mitochondrial metabolism (Courtney et al., 2018; Crooks et al., 2021; Mayr et al., 2008; Simonnet et al., 2003; Tomlinson et al., 2002). Decreases in CI expression and/or heteroplasmic mutations in mitochondrial DNA CI genes have also been reported in diverse cancers conditions, suggesting that activation of reductive mitochondrial anabolism may be a metabolic feature of those cancers (Gorelick et al., 2021; Reznik et al., 2017). On the contrary, CI inhibitors have also been identified as an actionable target for cancer treatment, which can decrease tumor growth by suppressing intratumoral aspartate (Baccelli et al., 2019; Gui et al., 2016; Martínez-Reyes et al., 2020; Molina et al., 2018; Sullivan et al., 2018; Wheaton et al., 2014). Notably, the responses to these mitochondrial inhibitors have been variable across clinical trials and cancer models suggesting that heterogeneity in cancer cell mitochondrial metabolism may govern the responses to CI inhibitors, and potentially to future inhibitors of reductive mitochondrial anabolism (Janku et al., 2021; Yap et al., 2019). Detailed understanding of mitochondrial metabolism phenotypes is thus critical for capitalizing on cancer cell metabolic changes to improve cancer therapy.

While our results demonstrate that decreased mitochondrial NAD+/NADH is a critical driver of aspartate synthesis in SDH-impaired cells, it remains unclear how these redox changes induce aspartate production. One likely contributor is pyruvate carboxylase, which can have increased expression and activity in SDH-mutant cancer cells (Cardaci et al., 2015; Lussey-Lepoutre et al., 2015). Another is reductive carboxylation of glutamine-derived alpha-ketoglutarate, whose relative contribution to aspartate production is increased in cells with defects in oxidative mitochondrial metabolism (Metallo et al., 2011; Mullen et al., 2012, 2014; Wise et al., 2011). Interestingly however, neither path directly consumes mitochondrial NADH, making the mechanistic connection between decreased mitochondrial NAD+/NADH and aspartate production in SDH-deficient cells an important goal for future study. Nonetheless, the observation that mito*LbNOX*, but not cyto*LbNOX*, blocks the benefits of CI inhibition in SDH-impaired cells localizes the site of effect to the mitochondria. These results therefore reveal that compartmentalized redox state can functionally alter cell phenotypes independent of the remainder of the cell and highlight that investigation of subcellular metabolic states will be essential to develop a complete understanding of cell metabolism.

## Acknowledgments

We thank members of the Sullivan lab and the Finley lab at MSKCC for discussion and feedback. This research was supported by the Proteomics & Metabolomics Shared Resource of the Fred Hutch/University of Washington Cancer Consortium (P30CA015704). M.L.H. was supported by the Molecular Medicine Training Program (NIH T32GM095421), a GO-MAP fellowship, and the NCI (R00CA218679-03S1). L.B.S. acknowledges support from the NCI (R00CA218679), the Andy Hill Cancer Research Endowment, and startup funds provided by the Fred Hutchinson Cancer Center. Support for L.B.S. and S.M.C. was also provided by a pilot award from the NCI Partnership for the Advancement of Cancer Research (U54CA132381).

## Contributions

M.L.H. and L.B.S. conceived of the project and performed all the experiments with assistance from E.Q., A.B.G.V., O.J.N., I.A.E., K.D., and P.H. S.M.C. performed GSEA on our proteomics dataset. L.B.S. supervised the project. M.L.H. and L.B.S. wrote the manuscript with input from all the authors.

## Competing Interests

None.

## Materials and Methods

### Cell Culture

Cell lines were acquired from ATCC (143B, HEK293T, 786-O, A549, HCT116), as a gift from Dr. W. Marston Linehan, NCI (UOK269), or as a gift from Dr. Stanley Lee, Fred Hutch (TF-1). Cell identities were confirmed by satellite tandem repeat profiling and cells were tested to be free from mycoplasma (MycoProbe, R&D Systems). Cells were maintained in Dulbecco’s Modified Eagle’s Medium (DMEM) (Gibco, 50-003-PB) supplemented with 3.7 g/L sodium bicarbonate (Sigma-Aldrich, S6297), 10% fetal bovine serum (FBS) (Gibco, 26140079) and 1% penicillin-streptomycin solution (Sigma-Aldrich, P4333). TF-1 cells were also supplemented with the 2 ng/ml Human Granulocyte Macrophage-Colony Stimulating Factor (GM-CSF) (Shenandoah Biotechnology, 100-08). Cells were incubated in a humidified incubator at 37°C with 5% CO_2_.

### Proliferation Assays

Cells were trypsinized (Corning, 25051CI), resuspended in media, and counted (Beckman Coulter Counter Multisizer 4 or Nexcelom Auto T4 Cellometer) and seeded overnight onto 6-well dishes (Corning, 3516) with an initial seeding density of 20,000 cells/well (143B, HEK293T, HCT116, 786-O, A549) or 30,000 cells/well (UOK269). After overnight incubation, 3-6 wells were counted for a starting cell count at the time of treatment. Cells were washed twice in phosphate-buffered saline (PBS) and 4 mL of treatment media was added. Experiments were conducted in DMEM without pyruvate (Corning 50-013-PB) supplemented with 3.7 g/L sodium bicarbonate 10% dialyzed fetal bovine serum (FBS) (Sigma-Aldrich, F0392) and 1% penicillin-streptomycin solution, with or without 1 mM sodium pyruvate (PYR) (Sigma-Aldrich, P8574), 1 mM 2-ketobutyric acid (AKB) (Sigma-Aldrich, K401), 1-20 mM Aspartic Acid (ASP) (Sigma-Aldrich, A7219), or 1 mM Asparagine (Sigma-Aldrich, A7094). Proliferation assays contain 1 mM PYR unless otherwise noted. Proliferation assays using Human Plasma-Like Medium (HPLM) (ThermoFisher, A4899101) were supplemented with 10% dialyzed FBS, 1% penicillin-streptomycin solution, with or without 20 mM ASP. Drug treatments included rotenone (Sigma-Aldrich, R8875), metformin (Sigma-Aldrich, D150959), piericidin A (Cayman Chemical, 15379), atpenin A5 (Cayman Chemical, 11898; AdipoGen, AG-CN2-0110; Abcam, ab144194; or Enzo Life Sciences, ALX-380-313), doxycycline hydrochloride (Sigma-Aldrich, D3447), AZD7545 (Cayman Chemical, 19282), antimycin A (Sigma-Aldrich, A8674), oligomycin (Sigma-Aldrich, 495455), and DMSO vehicle (D2650). Cells were incubated in a humidified incubator at 37°C with 5% CO_2_ then counted after 4-6 days, and proliferation rate was determined by the following equation: Proliferation rate (doublings per day, 1/d) = (log_2_(final cell count / initial cell count))/total days.

### Lentiviral Production and Infection

The following plasmids were obtained from DNASU Plasmid Repository: pLenti6.3-V5-DEST_SLC1A3, pLenti6.3-V5-DEST_SDHB, and pDONR201-NDI1. pLX304-gpASNase1 and pDONR-cyto*LbNOX* were previously described (Luengo et al., 2021; Sullivan et al., 2018). Cyto*LbNOX* was then cloned into pLX304 (Addgene, 25890) and NDI1 was cloned into pInducer20 (Addgene, 44012, gift from Stephen Elledge) using LR Clonase II (Fisher, 11791100). Lentivirus was generated by transfection of HEK293T cells with expression construct plasmid DNA with pMDLg/pRRE (Addgene, 12251), pRSV-Rev, (Addgene, 12253) and pMD2.G (Addgene, 12259) packaging plasmids. and FuGENE transfection reagent (Fisher, PRE2693) in DMEM (Fisher, MT10017CV) without FBS or penicillin-streptomycin. The supernatant containing lentiviral particles was filtered through 0.45 µM membrane (Fisher, 9720514) and was supplemented with 8 µg/µL polybrene (Sigma, TR-1003-G) prior to infection. 143B and UOK269 cells were seeded at 50% confluency in 6 well dishes and centrifuged with lentivirus (900 xg, 90 mins, 30°C). After 24 hours media was replaced with fresh media and after 48 hours cells were treated with either 1 µg/mL blasticidin (Fisher, R21001) or 1 mg/mL G418 (Sigma, A1720) and maintained in selection media until all uninfected control cells died.

### Generation of mitoLbNOX expressing cells

*LbNOX*-FLAG DNA was amplified from pDONR-cyto*LbNOX* by PCR, an oligonucleotide containing the zmLOC100282174 mitochondrial translocation sequence (MTS), which causes potent mitochondrial localization, (Chin et al., 2018) was purchased (Integrated DNA Technologies) and the pENTR1A vector (Fisher, A10462) was amplified by PCR to create a linear fragment. The three fragments were assembled using the NEBuilder HiFi DNA Assembly Cloning Kit (New England BioLabs, E2621) to generate pENTR1A-mito*LbNOX*, which was then used as a donor to transfer mitoLbNOX into pLenti-CMV-Hygro-DEST (w117-1) (Addgene, 17454 a gift from Eric Campeau & Paul Kaufman) using Gateway LR Clonase II (Fisher, 11791020). Lentivirus was generated with pLentiCMV-mito*LbNOX* as described above and 143B cells were infected and then selected in 150 µg/mL Hygromycin B (Sigma, H7772) for four days.

*LbNOX*-FLAG primers:

5’-GCCGCGAGACCGTATGCTCATAAGGTCACCGTG-3’ (fw primer)

5’-CAAGAAAGCTGGGTCTAGTTACTTGTCATCGTCATCC-3’ (rv primer)

pENTR primers:

5’-CTAGACCCAGCTTTCTTGTAC-3’ (fw)

5’-CTGGCTTTTAGTAAGCGAATTC-3’ (rv)

MTS sequence:

5’-CGCTTACTAAAAGCCAGGCCACCATGGCACTGCTTCGCGCCGCCGTTTCAGAACTCAGACGGAGAGGACGGGGTGCGCTTACTCCCCTCCCGGCGCTGTCTAGCTTGCTTTCCTCACTTAGCCCCCGAAGTCCCGCCTCAACGCGCCCAGAGCCAAACAATCCACACGCAGATCGACGCCATGTCATCGCTTTGAGGCGATGCCCCCCACTTCCTGCCTCTGCCGTTCTGGCACCTGAACTCCTGCATGCACGAGGATTGCTCCCGAGACATTGGTCTCATGCCTCTCCCTTGTCCACGTCCTCTTCATCCAGTAGACCAGCAGATAAGGCGCAGTTGACCTGGGTCGATAAATGGATCCCAGAAGCCGCGAGACCGTAT-3’

### Generation of Knockout Cell Lines

This protocol was adapted from Hoellerbauer et al., 2020. Three chemically synthesized 2’-O-methyl 3’phosphorothioate-modified single guide RNA (sgRNA) sequences targeting SDHB were purchased (Synthego). The sgRNA sequences were 5’-UCGCCCUCUCCUUGAGGCG-3’, ‘5’-AGAAAUUUGCCAUCUAUCGA-3’, and 5’-CUUUGUUAGAUGUGGCCCCA-3’. Each sgRNA was resuspended in nuclease-free water, combined with SF buffer (Lonza, V4XC-2032), and sNLS-spCas9 (Aldevron, 9212). 2×10^5^ 143B or HEK293T cells were resuspended in the resulting solution containing ribonucleoprotein complexes (RNPs) and electroporated using a 4D-Nucleofector (Amaxa, Lonza) programs FP-133 (143B) and DS-150 (HEK293T). Nucleofected cells were then moved to a 12-well plate (Corning, 3513) and, after achieving confluence, were single-cell cloned by limiting dilution by plating 0.5 cells/well in a 96 well plate. SDHB gene knockout was confirmed using western blots on the nucleofected pool and each single cell clone used in this study.

### Oxygen Consumption

Oxygen consumption measurements were conducted using an Agilent Seahorse Xfp Analyzer. 143B or UOK269 cell lines were trypsinized and seeded at 2.5×10^5^ cells/well of a Seahorse XFp cell culture miniplate (Agilent, 103025-1000) overnight in 80 µL media. The following day, 100 µL cell culture media was added with or without pretreatment, as indicated. Before the assay Seahorse XFp sensor cartridges (Agilent, 103022-100) were incubated with calibrant solution, following manufacturer’s instructions and loaded with injection solutions yielding the following final concentrations: AA5; 5 µM, rotenone; 100 nM. AZD7545; 5 µM, antimycin A; 10 µM. Following each assay, cells of each well were counted by Coulter Counter and mitochondrial oxygen consumption rate was determined as the measured oxygen consumption rate minus the average of post-antimycin oxygen consumption rates, per 100,000 cells.

### Mitochondrial Fractionation

2×10^7^ cells were trypsinized, washed with PBS, and centrifuged (300 xg, 5 min, 4°C). Pellets were washed with ice-cold PBS and resuspended in Homogenization buffer (10 mM Tris-HCl pH 6.7, 10 mM KCl, 0.15 mM MgCl_2_, 1 mM PMSF (Sigma, 10837091001)). Cells were then vortexed, incubated on ice for 2 minutes, then transferred to a Dounce Homogenizer (Sigma, D9063) and homogenized with 40-60 strokes. Cells were transferred to mitochondrial suspension buffer (10 mM Tris-HCl pH 6.7, 0.15 mM MgCl_2_, 0.25 mM sucrose, 1 mM PSMF) and centrifuged (700 xg, 10 min, 4°C). The supernatant was transferred and centrifuged again (10,500 xg, 15 min, 4°C). The resulting supernatant (cytosolic fraction) was transferred and centrifuged once more, (17,000 xg, 15 min, 4°C) while the pellet containing the mitochondrial fraction was washed in suspension buffer and centrifuged again (12,000 xg, 15 min, 4°C). Mitochondrial proteins were then harvested with RIPA buffer (see below).

### Western Blotting

Protein lysates were harvested in RIPA buffer (Sigma, R0278) supplemented with protease inhibitors (Fisher, A32953). Protein concentration was determined using a Bicinchoninic Acid Assay (Fisher, 23225) using BSA as a standard. Equal amounts of protein were denatured with Bolt 4x Loading Dye (Thermo, B0007) and Bolt 10x reducing agent (Fisher, B0004), heated at 95°C for 5 min, and loaded onto 4-12% by SDS-polyacrylamide gels (Fisher, NW04127). After electrophoretic separation, proteins were transferred onto a 0.22 mm nitrocellulose using iBlot2 transfer stacks (Fisher, IB23001) and transferred with the P0 system setting. Membranes were blocked with 5% milk in Tris-buffered saline with 0.1% Tween-20 (TBS-T) and incubated at 4°C overnight with the following antibodies: anti-FLAG (Sigma, F1804; 1:1,000), anti-SLC1A3 (Genetex, GTX20262; 1:500), anti-SDHB (Atlas, HPA002868; 1:1,000), anti-NDUFA8 (Atlas, HPA041510; 1:1,000), anti-Vinculin (Sigma, SAB4200729; 1:10,000), and anti-Tubulin (Sigma, T6199; 1:10,000). The next morning, membranes were washed three times with TBS-T and the following secondary antibodies were added: 800CW Goat anti-Mouse IgG (LiCOR, 926-32210; 1:15,000), 680RD Goat anti-Rabbit IgG (LiCOR, 926-68071; 1:15,1000). Membranes were washed three more times with TBS-T and imaged on a LiCOR Odyssey Near-Infrared imaging system.

### TMT-Quantitative Mitochondrial Proteomics

*<u>Sample preparation</u>*. Mitochondrial fractions of WT 143B, SDHB AB 143B, EP SDHB KO 143B, and LP SDHB KO 143B cells in quadruplicate were extracted using the protocol described above. Mitochondrial pellets were kept at -80°C until analysis. *<u>Disulfide bond reduction/alkylation</u>*. Protein solutions (100 μg) were diluted to 2 μg/μL in 100 mM ammonium bicarbonate. Protein disulfide bonds were reduced by adding tris (2-carboxyethyl) phosphine to a final concentration of 5 mM and mixing at room temperature for 15 min. The reduced proteins were alkylated by adding 2-chloroacetamide to a final concentration of 10 mM and mixing in the dark at room temperature for 30 min. Excess 2-chloroacetamide was quenched by the addition of dithiothreitol to 10 mM and mixing at room temperature for 15 min. *<u>Methanol-chloroform precipitation and protease digestion</u>*. Samples (100 μg) were diluted to 1 μg/μL with 100 mM ammonium bicarbonate in a 1.5 mL Eppendorf low bind tube. Protein precipitation was done as follows: 400 μL of methanol was added to each sample and vortexed for 5 seconds. 100 μL of chloroform was added to each sample and vortexed for 5 seconds. 300 μL of water was added to each sample and vortexed for 5 seconds. The samples were centrifuged for 1 min at 14,000 xg. The aqueous and organic phases were removed, leaving a protein wafer in the tube. The protein wafers were washed with 400 μL of methanol and centrifuged at 21,000 g at room temperature for 2 min. The supernatants were removed, and the pellets were allowed to air dry, but not to complete dryness. The samples were resuspended in 70 μL 100 mM HEPES (pH 8.5) and digested with rLys-C protease (100:1, protein to protease) with mixing at 37 °C for 4 hr. Trypsin protease (100:1, protein to protease) was added and the reaction was mixed overnight at 37 °C. *<u>TMT-labeling</u>*. TMTpro16plex labeling reagent (Pierce) (500 μg) was brought up in 30 μL acetonitrile and added to the digested peptide solution (100 μg) yielding a final organic concentration of 30% (v/v) and mixed at room temperature for 1 hr. A 2 μg aliquot from each sample was combined, dried to remove the acetonitrile, concentrated on a C18 ZipTip (Millipore) and analyzed via LC/MS as a “label check”. The label check was used for two purposes. First, it ensured the TMT labeling efficiency was greater than 95%. Second, sample equalization volumes were determined after summing the total intensity from each sample TMT “channel” and calculating the volume of each sample that should be mixed to provide equal amounts from each sample in the final mixture. After labeling efficiency was determined, the reactions were quenched with hydroxylamine to a final concentration of 0.3% (v/v) for 15 min with mixing. The TMTpro16plex labeled samples were pooled at 1:1 ratio based on calculated equalization volumes and concentrated by vacuum centrifugation to remove acetonitrile. Half of the material was desalted over an Oasis HLB 3cc cartridge (Waters) and taken to dryness. *<u>bRP Fractionation</u>*. Pierce’s High pH Reversed-Phased Peptide Fractionation kit (part# 84868) was used to fractionate the sample (100 ug) into 8 fractions with cuts at 5, 7.5, 10.0, 12.5, 15, 17.5, 20.0, 22.5, 25.0 and 50.0% acetonitrile in 0.1% triethylamine following the manufacturer’s protocol. The fractions were dried in a speedvac. *<u>Mass Spectrometry Analysis</u>*. The generated basic reverse phase fractions were brought up in 2% acetonitrile in 0.1% formic acid (20 μL) and analyzed (2 μL) by LC/ESI MS/MS with a Thermo Scientific Easy1200 nLC (Thermo Scientific, Waltham, MA) coupled to a tribrid Orbitrap Eclipse with FAIMS pro (Thermo Scientific, Waltham, MA) mass spectrometer. In-line de-salting was accomplished using a reversed-phase trap column (100 μm × 20 mm) packed with Magic C_18_AQ (5-μm, 200 Å resin; Michrom Bioresources, Auburn, CA) followed by peptide separations on a reversed-phase column (75 μm × 270 mm) packed with ReproSil-Pur C_18_AQ (3-μm, 120 Å resin; Dr. Maisch, Baden-Würtemburg, Germany) directly mounted on the electrospray ion source. A 180-minute gradient from 4% to 44% B (80% acetonitrile in 0.1% formic acid/water) at a flow rate of 300 nL/minute was used for chromatographic separations. A spray voltage of 2300 V was applied to the electrospray tip in-line with a FAIMS pro source using varied compensation voltage -40, -60, -80 while the Orbitrap Eclipse instrument was operated in the data-dependent mode. MS survey scans were in the Orbitrap (Normalized AGC target value 300%, resolution 120,000, and max injection time 50 ms) with a 3 sec cycle time and MS/MS spectra acquisition were detected in the linear ion trap (Normalized AGC target value of 100% and injection time 50 ms) using CID activation with a normalized collision energy (NCE) of 32% using turbo speed scan. Selected ions were dynamically excluded for 60 seconds after a repeat count of 1. Following MS2 acquisition, real time searching (RTS) was employed, and spectra were searched against a Human database (UP00005640 Human 120119) using COMET. Searches were performed with settings for the proteolytic enzyme trypsin. Maximum missed cleavages were set to 1 and maximum variable modifications on peptides was set to 3. Variable modifications included oxidation (+15.995 Da on M) with static modifications TMTpro (+304.207 DA on K) and carbamidomethyl (+57.021 on C). Maximum search time was 35 ms. Scoring thresholds were set to the following: Xcorr 1.4, dCn 0.1, precursor PPM 10 and charge state 2. TMT Synchronous precursor selection (SPS) MS3 was collected on the top 10 most intense ions detected in the MS2 spectrum. SPS-MS3 precursors were subjected to higher energy collision-induced dissociation (HCD) for fragmentation with an NCE of 45% and analyzed using the Orbitrap (Normalized AGC target value of 400%, resolution 60,000 and max injection time 118 ms). *<u>Data Analysis</u>*. Data analysis was performed using Proteome Discoverer 2.5 (Thermo Scientific, San Jose, CA). The data were searched against a Human database (UP00005640 Human 120119) that included common contaminants (cRAPome). Searches were performed with settings for the proteolytic enzyme trypsin. Maximum missed cleavages were set to 2. The precursor ion tolerance was set to 10 ppm and the fragment ion tolerance was set to 0.6 Da. Dynamic peptide modifications included oxidation (+15.995 Da on M). Dynamic modifications on the protein terminus included acetyl (+42.-11 Da on N-terminus), Met-loss (−131.040 Da on M) and Met-loss+Acetyl (−89.030 Da on M) and static modifications TMTpro (+304.207 Da on any N-termius), TMTpro (+304.207 DA on K) and carbamidomethyl (+57.021 on C). Sequest HT was used for database searching. All search results were run through Percolator for scoring.

### Polar Metabolite Extractions

*<u>Adherent cells</u>*. For standard metabolic analysis, cells were seeded overnight at 2×10^5^ cells per well of a 6-well dish. The next morning, cells were washed twice with PBS and changed to the indicated medias supplemented with 10% dialyzed FBS, 1% penicillin-streptomycin, and treatments as indicated, and returned to the tissue culture incubator. After 6 hours, polar metabolites were extracted from cells by three washes with ice-cold blood bank saline, (Fisher, 23293184) then 250 µL of 80% HPLC grade methanol in HPLC grade water containing 2.5 µM D8 Valine was added to each well and cells were scraped with the back of a P1000 pipet tip and transferred to Eppendorf tubes. Tubes were centrifuged (17,000 xg, 15 mins, 4°C) and the supernatant containing polar metabolites was transferred to a new centrifuge tube and placed in a centrivap until lyophilized. *<u>Suspension cells</u>*. TF-1 cells were seeded at 2.5×10^5^ cells per well of a 6-well dish the night before treatment and extraction. The next day, cells were washed with PBS and treated with media containing 2 ng/mL human GM-CSF, 10% dialyzed FBS, and the treatments as indicated. After 6 hours, cells were collected and washed in 1 mL of ice-cold saline containing 2% dialyzed FBS. Metabolism was quenched and metabolites were extracted with 600 µL of 80% HPLC grade methanol in HPLC grade water. Samples were centrifuged, (17,000 xg, 15 mins, 4°C) moved to new Eppendorf tubes, lyophilized, and stored at -20°C until analysis.

### Isotope Tracing

*<u>Aspartate Uptake</u>*. SLC1A3 143B cells were plated at 2×10^5^ cells per well of a 6-well dish. The next morning, cells were washed twice with PBS and changed to DMEM without pyruvate supplemented with 10% dialyzed FBS, 1% penicillin-streptomycin, and treated with the indicated concentrations of 1,4-^13^C2 aspartate (Cambridge Isotopes, CLM-4455) for one hour. *<u>Glutamine Tracing</u>*. WT 143B cells were seeded at 2×10^5^ cells and UOK269 cell lines were plated at 3×10^5^ cells per well of a 6-well dish. The next morning, cells were washed twice with PBS and changed to DMEM without glucose, glutamine, pyruvate, or phenol red (Sigma, D5030) supplemented with 10% dialyzed FBS, 1% penicillin-streptomycin, 1 mM pyruvate, 25 mM ^12^C glucose (Sigma, G7528), and 4 mM U-^13^C glutamine (Cambridge Isotopes, CLM-1822). 143B cells were treated as indicated for 6 hours. *<u>Glucose Tracing</u>*. WT 143B cells were seeded at 2×10^5^ cells per well of a 6-well dish. The next morning, cells were washed twice with PBS and swapped to DMEM without glucose, glutamine, pyruvate, or phenol red supplemented with 10% dialyzed FBS, 1% penicillin-streptomycin, 1 mM AKB, 25 mM U-^13^C glucose (Cambridge Isotopes, CLM-1396), and 4 mM ^12^C glutamine (Sigma, G5792). Polar metabolites were extracted with the above technique.

### Liquid Chromatography-Mass Spectrometry (LCMS)

Lyophilized samples were resuspended in 80% HPLC grade methanol in HPLC grade water and transferred to liquid chromatography-mass spectrometry (LCMS) vials for measurement by LCMS. Metabolite quantitation was performed using a Q Exactive HF-X Hybrid Quadrupole-Orbitrap Mass Spectrometer equipped with an Ion Max API source and H-ESI II probe, coupled to a Vanquish Flex Binary UHPLC system (Thermo Scientific). Mass calibrations were completed at a minimum of every 5 days in both the positive and negative polarity modes using LTQ Velos ESI Calibration Solution (Pierce). Polar Samples were chromatographically separated by injecting a sample volume of 1 μL into a SeQuant ZIC-pHILIC Polymeric column (2.1 × 150 mm 5 mM, EMD Millipore). The flow rate was set to 150 mL/min, autosampler temperature set to 10 °C, and column temperature set to 30 °C. Mobile Phase A consisted of 20 mM ammonium carbonate and 0.1 % (v/v) ammonium hydroxide, and Mobile Phase B consisted of 100 % acetonitrile. The sample was gradient eluted (%B) from the column as follows: 0-20 min.: linear gradient from 85 % to 20 % B; 20-24 min.: hold at 20 % B; 24-24.5 min.: linear gradient from 20 % to 85 % B; 24.5 min.-end: hold at 85 % B until equilibrated with ten column volumes. Mobile Phase was directed into the ion source with the following parameters: sheath gas = 45, auxiliary gas = 15, sweep gas = 2, spray voltage = 2.9 kV in the negative mode or 3.5 kV in the positive mode, capillary temperature = 300 °C, RF level = 40 %, auxiliary gas heater temperature = 325 °C. Mass detection was conducted with a resolution of 240,000 in full scan mode, with an AGC target of 3,000,000 and maximum injection time of 250 msec. Metabolites were detected over a mass range of 70-1050 *m/z*. Quantitation of all metabolites was performed using Tracefinder 4.1 (Thermo Scientific) referencing an in-house metabolite standards library using ≤ 5 ppm mass error. Data from stable isotope labeling experiments includes correction for natural isotope abundance using IcoCor software v.2.2.

### Complex I Activity Assay

Complex I activity was measured using an assay kit (Abcam, ab109721) according to the manufacturer’s instructions. Absorbances at 450 nm were read every 30 seconds for 40 minutes on a Tecan Infinite M Plex microplate reader.

### Gene Set Enrichment Analysis

Gene Set Enrichment Analysis (GSEA) (Subramanian et al., 2005) was conducted on the proteomics dataset utilizing 149 hierarchical, mitochondrial specific pathways curated by MitoCarta3.0 (Rath et al., 2021) as the *a priori* defined gene/protein sets. Proteins were ranked by the signed (negative or positive by the direction of the fold-change), -log of the adjusted p-value, both of which were calculated during differential expression analysis between LP and WT cells, and used as input for GSEA. Individual MitoCarta3.0 pathways were evaluated for enrichment of upregulated proteins (positive normalized enrichment score) and downregulated proteins (negative normalized enrichment score). GSEA analysis was implemented in R (*R Core Team 2020*) (version 4.0.3) using the Bioconductor package fgsea (Korotkevich et al., 2021). Adjusted p-values and normalized enrichment scores have been reported.

### Statistical Analysis

All graphs and statistical analyses were made in GraphPad Prism 9.0. Technical replicates, defined as parallel biological samples independently treated, collected, and analyzed during the same experiment, are shown. Experiments were verified with ≥ 2 independent repetitions showing qualitatively similar results. Details pertaining to all statistical tests can be found in the figure legends.

## Notes

### Competing Interest Statement

The authors have declared no competing interest.

