## Supplemental Figures for "Mitochondrial Redox Adaptations Enable Aspartate Synthesis in SDH-deficient Cells"

### Supplemental Figure 1.

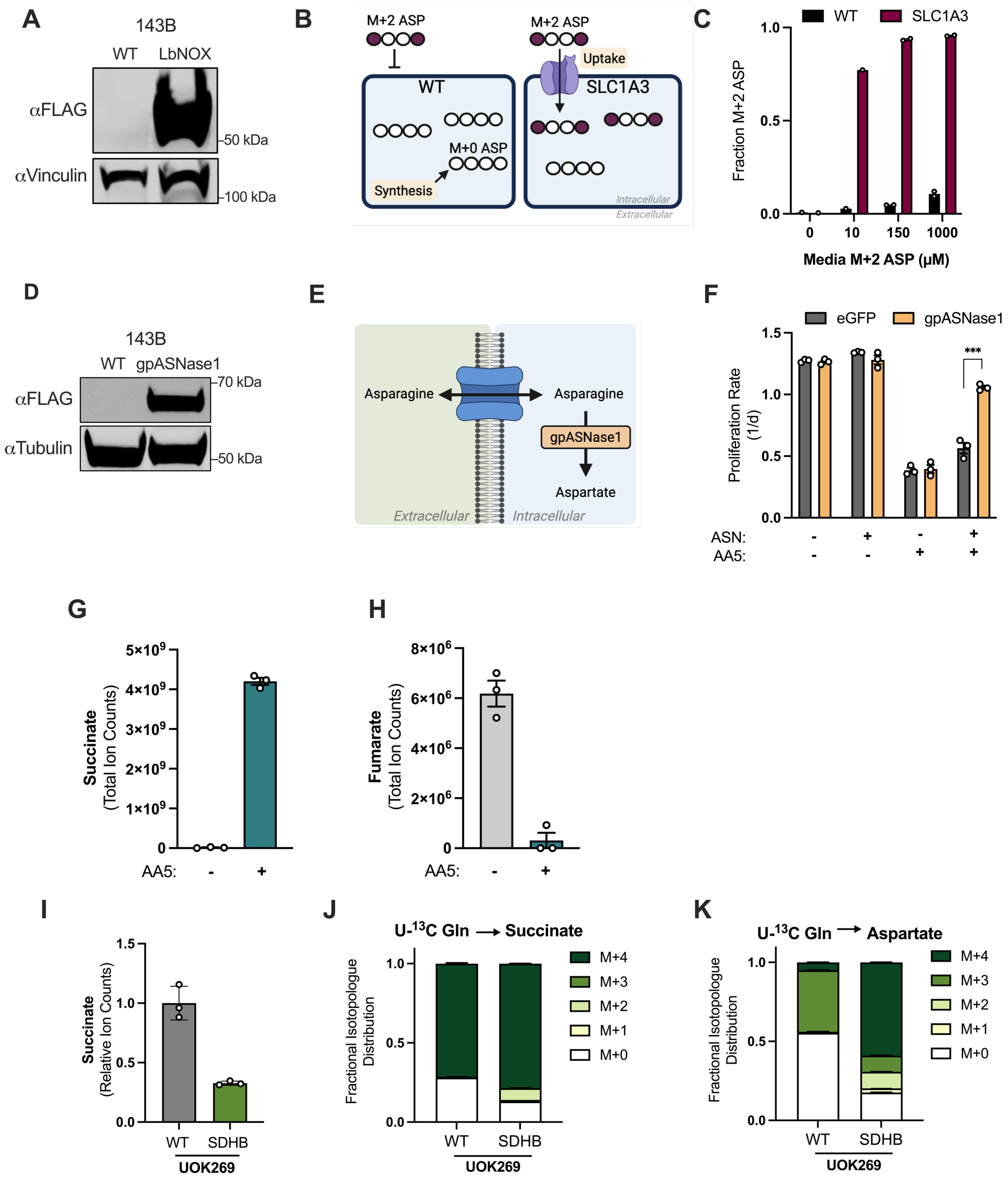

**Supplemental Figure 1. Alternative methods of aspartate acquisition and characterization of metabolic phenotypes in SDH-impaired cells.**

(A) Western blot for FLAG and vinculin from WT 143B cells and 143B cells expressing FLAG-tagged cyto*LbNOX*. Vinculin is used as a loading control.
(B) Schematic demonstrating how the ASP transporter SLC1A3 allows for cells to uptake aspartate, which can be measured by the incorporation of isotopically labeled extracellular aspartate (M+2).
(C) Fractional labeling of aspartate measured by LCMS metabolomics from 143B cells with or without SLC1A3 expression after 1 hour of exposure to the indicated concentrations of 1,4-13C labeled (M+2) aspartate (n=1 or n=2).
(D) Western blot for FLAG and tubulin from WT 143B cells and 143B cells expressing FLAG-tagged gpASNase1. Tubulin is used as a loading control.
(E) Schematic depicting how expression of gpASNase1 permits environmental asparagine (ASN) to be used to support intracellular aspartate levels.
(F) Proliferation rates of 143B cells expressing eGFP or gpASNase1 cultured in DMEM and supplemented with vehicle ($H_2O$) or 1 mM asparagine (ASN) and treated with vehicle (DMSO) or 5 µM AA5 (n=3).
(G) Succinate levels measured by LCMS metabolomics from 143B cells treated with vehicle (DMSO) or 5 µM AA5 for 6 hours (n=3).
(H) Fumarate levels measured by LCMS metabolomics from 143B cells treated with vehicle (DMSO) or 5 µM AA5 for 6 hours (n=3).
(I) Relative succinate levels measured by LCMS metabolomics from WT and SDHB UOK269 cells (n=3).
(J) Fractional isotopologue distribution of succinate measured by LCMS metabolomics of WT or SDHB UOK269 cells cultured in glutamine free DMEM supplemented with 4 mM U-$^{13}$C glutamine for 6 hours (n=3).
(K) Fractional isotopologue distribution of aspartate measured by LCMS metabolomics of WT or SDHB UOK269 cells cultured in glutamine free DMEM supplemented with 4 mM U-$^{13}$C glutamine for 6 hours (n=3).

Data are plotted as means ± standard deviation (SD) and compared with an unpaired two-tailed student's t-test. p<0.05*, p<0.01**, p<0.00***, p<0.0001****

### Supplemental Figure 2.

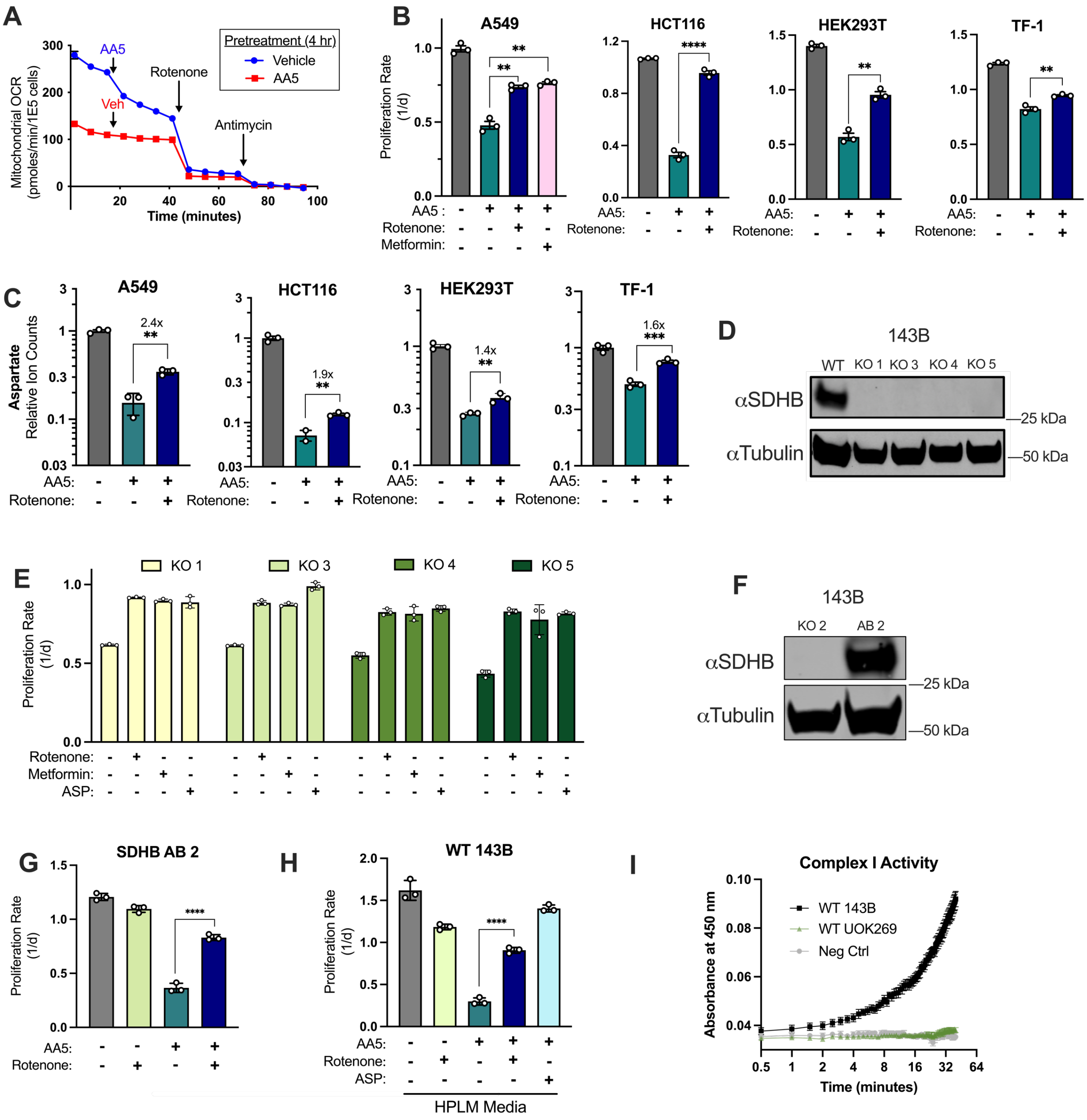

**Supplemental Figure 2. Characterization of interactions between complex I inhibition and SDH status.**

(A) Mitochondrial oxygen consumption rates in WT 143B cells pre-treated with vehicle (DMSO) or 5 μM AA5 for 4 hours, then injected with vehicle (DMSO) or 5 μM AA5 as indicated, followed by treatment of all conditions with 100 nM rotenone and 10 μM antimycin (n=3).
(B) Proliferation rates of A549, HCT116, HEK293T, and TF-1 cells cultured in DMEM and treated with vehicle (DMSO), AA5, or AA5 and rotenone. A549 cells were treated with 2.5 μM AA5, 80 nM rotenone, and 500 μM metformin and HCT116, HEK293T, and TF-1 cells were treated with 5 μM AA5 and 50 nM rotenone (n=3).
(C) Relative aspartate levels measured by LCMS metabolomics of A549, HCT116, HEK293T, and TF-1 cells cultured in DMEM and treated with vehicle (DMSO), AA5, or AA5 and rotenone for 6 hours as done in B (n=3).
(D) Western blot for SDHB and tubulin from WT 143B cells and four SDHB KO 143B clones. Tubulin is used as a loading control.
(E) Proliferation rates of corresponding SDHB KO clones from D treated with vehicle (DMSO), 50 nM rotenone, 1 mM metformin, or 20 mM ASP (n=3).
(F) Western blot for SDHB and tubulin from SDHB KO clone 2 (KO 2) and the cells with SDHB cDNA added back (AB 2). Tubulin was used as a loading control.
(G) Proliferation rates of SDHB AB 2 cells treated with vehicle (DMSO), 50 nM rotenone, 5 μM AA5, or 5 μM AA5 and 50 nM rotenone (n=3).
(H) Proliferation rates of WT 143B cells cultured in HPLM and treated vehicle (DMSO), 50 nM rotenone, 5 μM AA5, 5 μM AA5 and 50 nM rotenone, or 5 μM AA5 and 20 mM ASP (n=3).
(I) Complex I activity assay (Abcam) of WT 143Bs and WT UOK269 cells (n=3).

Fold change is indicated above brackets in C. Data are plotted as means ± standard deviation (SD) except in A and I which are means ± standard error of the mean (SEM) and compared with an unpaired two-tailed student's t-test. $p<0.05$*, $p<0.01$**, $p<0.00$***, $p<0.0001$****

**Supplemental Figure 3.**

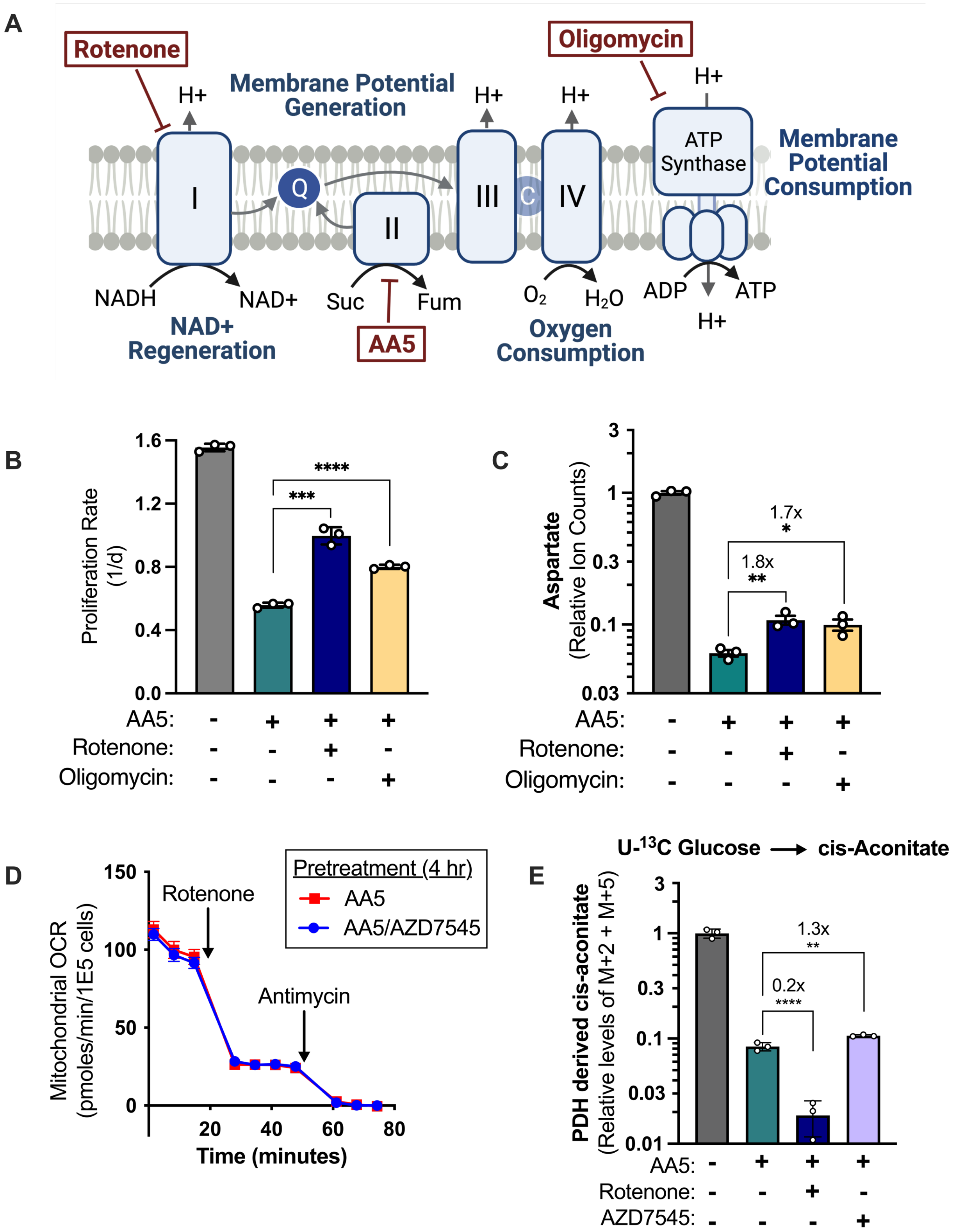

**Supplemental Figure 3. Effects of ETC and PDK inhibition in SDH-impaired cells.**

(A) Schematic showing the metabolic roles of the electron transport chain and ATP synthase and depicting the sites of action for rotenone, AA5, and oligomycin.
(B) Proliferation rates of WT 143B cells cultured in DMEM and treated with Vehicle (DMSO), 5 μM AA5, 5 μM AA5 and 50 nM rotenone, or 5 μM AA5 and 1 μM oligomycin (n=3).
(C) Aspartate levels of WT 143B cells cultured in DMEM and treated with Vehicle (DMSO), 5 μM AA5, 5 μM AA5 and 50 nM rotenone, or 5 μM AA5 and 1 μM oligomycin for 6 hours (n=3).
(D) Mitochondrial oxygen consumption rates of WT 143B cells pre-treated for 4 hours with 5 μM AA5 or 5 μM AA5 and 5 μM AZD7545 with indicated injections of 100 nM rotenone and 10 μM antimycin (n=3).
(E) Combined M+2 and M+5 cis-aconitate levels relative to vehicle treated cells measured by LCMS metabolomics from WT 143B cells cultured in glucose/pyruvate free DMEM supplemented with 1 mM AKB and 25 mM U-$^{13}$C glucose and treated with vehicle (DMSO), 5 μM AA5, 5 μM AA5 and 50 nM rotenone, or 5 μM AA5 and 5 μM AZD7545 for 6 hours (n=3).

Fold change is indicated above brackets in C and E. Data are plotted as means ± standard deviation (SD) except in D which are means ± standard error of the mean (SEM) and compared with an unpaired two-tailed student's t-test. p<0.05*, p<0.01**, p<0.00***, p<0.0001****

### Supplemental Figure 4.

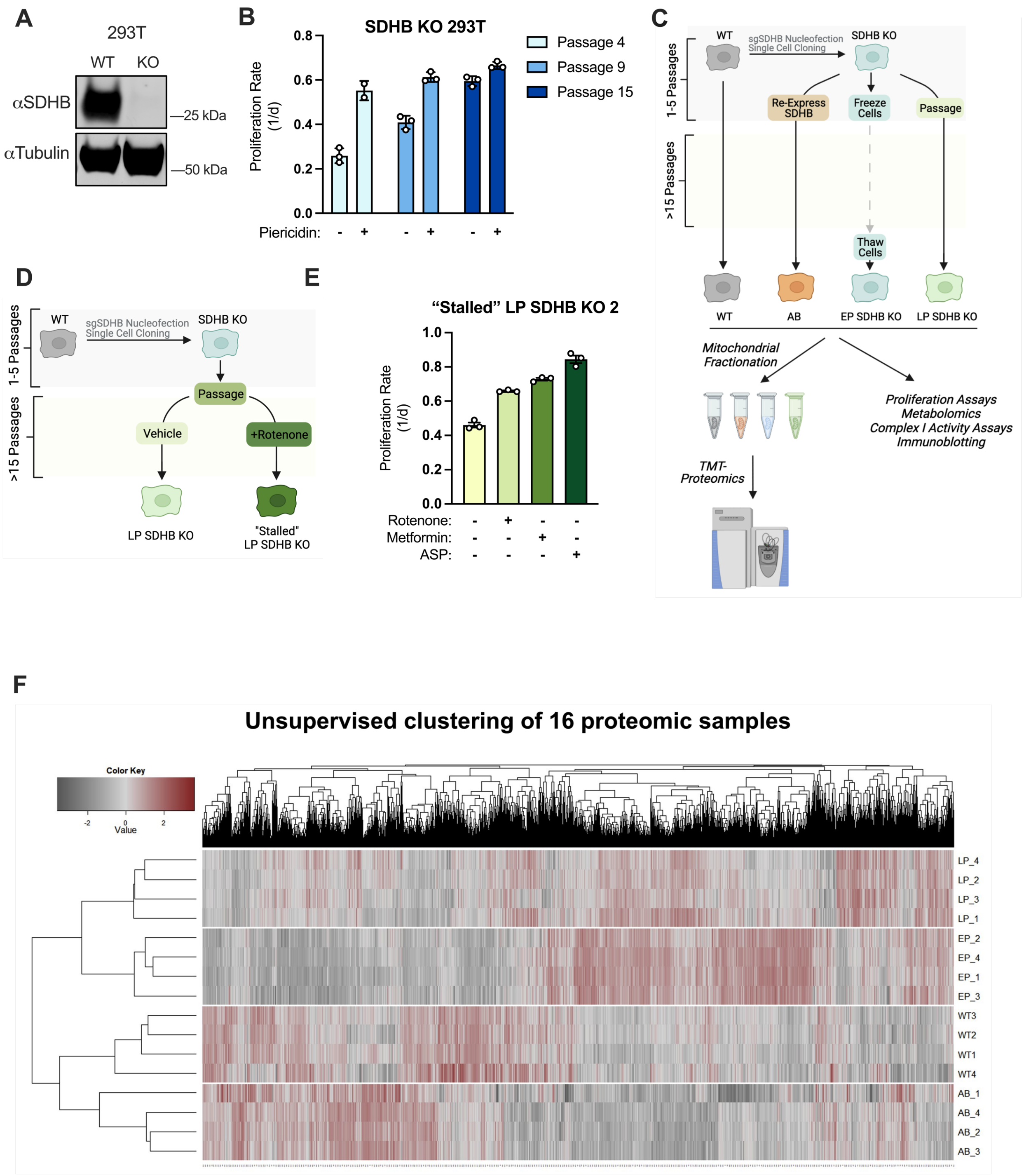

**Supplemental Figure 4. Characterization of adaptations in SDHB KO cells.**

(A) Western blot for SDHB and tubulin from WT and SDHB KO HEK293T cells. Tubulin is used as a loading control.
(B) Proliferation rates of SDHB KO HEK293T cells cultured in DMEM from different passage numbers. For each passage, cells were treated vehicle (DMSO) or 1 μM piericidin A, an inhibitor of CI.
(C) Schematic showing how SDHB KO 143B clones were used to generate SDHB addback cells (AB), early passage SDHB KO cells (EP), or late passage (LP) SDHB KO cells. The diagram also depicts how samples were acquired for mitochondrial proteomics and other experiments.
(D) Schematic illustrating how SDHB KO 143B cells were maintained in culture with or without continuous rotenone to achieve late passage (LP) status or “Stalled” LP status.
(E) Proliferation rates of “Stalled” LP SDHB KO 143B cells treated with vehicle (DMSO), 50 nM rotenone, 1 mM metformin, or 20 mM ASP.
(F) Unsupervised clustering from mitochondrial proteomics data of the four cell lines listed in C (n=4).

Data are plotted as means ± standard deviation (SD) and compared with an unpaired two-tailed student’s t-test. $p<0.05$*, $p<0.01$**, $p<0.00$***, $p<0.0001$****
